# Brain peak width of skeletonised mean diffusivity (PSMD), processing speed, and other cognitive domains

**DOI:** 10.1101/385013

**Authors:** Ian J. Deary, Stuart J. Ritchie, Susana Muñoz Maniega, Simon R. Cox, Maria C. Valdés Hernández, John M. Starr, Joanna M. Wardlaw, Mark E. Bastin

**Affiliations:** Department of Psychology, University of Edinburgh, Edinburgh, United Kingdom EH8 9JZ; Centre for Cognitive Ageing and Cognitive Epidemiology, University of Edinburgh, Edinburgh, United Kingdom EH8 9JZ; Alzheimer Scotland Dementia Research Centre, University of Edinburgh, Edinburgh, United Kingdom EH8 9JZ; Brain Research Imaging Centre, Division of Neuroimaging Sciences, University of Edinburgh, Edinburgh, United Kingdom EH16 4SB; Scottish Imaging Network, A Platform for Scientific Excellence (SINAPSE), University of Edinburgh, Edinburgh, United Kingdom EH16 4SB; Edinburgh Dementia Research Centre, Dementia Research Institute, University of Edinburgh, Edinburgh, United Kingdom EH16 4SB

**Keywords:** ageing, cognition, processing speed, structural MRI, diffusion MRI, white matter, PSMD

## Abstract

It is suggested that the brain’s peak width of skeletonised water mean diffusivity (PSMD) is a neuro-biomarker of processing speed, a crucial contributor to cognitive ageing. We tested whether PSMD is more strongly correlated with processing speed than with other cognitive domains, and more strongly than other structural brain MRI indices. Participants were 731 Lothian Birth Cohort 1936 members, mean age 73 years (SD=0.7); analytical sample was 656-680. Cognitive domains tested were: processing speed (5 tests), visuospatial (3), memory (3), and verbal (3). Brain-imaging variables included PSMD, white matter diffusion parameters and hyperintensity volumes, grey and white matter volumes, and perivascular spaces. PSMD was significantly associated with all processing speed tests; absolute standardised beta values were 0.11 to 0.23 (mean = 0.17). Other structural brain-imaging variables correlated as or more strongly. PSMD was significantly associated with processing speed (−0.27), visuospatial (−0.23), memory (−0.17), and general cognitive ability (−0.25). PSMD correlated with processing speed: but not more strongly than with other cognitive domains; and not more strongly than other brain-imaging measures.

## 1. Introduction

Cognitive functions such as processing speed, reasoning, and some aspects of memory decline, on average, as people grow older, with deleterious effects on people’s quality of life (Tucker-Drob, 2011; Salthouse, 2017; Institute of Medicine, 2015). Processing speed has a special place among the cognitive domains; it has been suggested as a foundation for long-standing differences in, and ageing of, other cognitive domains (Salthouse, 1996, 2000; Deary, 2000; Verhaeghen, 2014; Ritchie et al., 2014). With regard to understanding cognitive ageing, detailed attention has been advocated in the study of the psychometric aspects of processing speed and its potential neurobiological correlates (Salthouse and Madden, 2008). Therefore, understanding the neurobiological foundations of people’s differences in, and ageing of, processing speed and other cognitive domains is a research priority (Penke et al., 2012; Salthouse, 2000).

Even among those with no overt disease, brain imaging-derived biomarkers of brain deterioration correlate with performance on cognitive tests; such markers include cerebral tissue volumes and brain atrophy, and indicators of white matter health and so-called cerebral small vessel disease (Arvanitakis et al., 2016; Cremers et al., 2016; Dong et al., 2015; Gazes et al., 2016; Liu et al., 2017; Salthouse et al., 2015). Cerebral small vessel disease is an important contributor to vascular-based cognitive deterioration (Shi and Wardlaw, 2016). Recognising that, a brain imaging marker named ‘peak width of skeletonized [water] mean diffusivity’ (PSMD) was developed and studied with respect to its association with processing speed (Baykara et al., 2016). It is based on skeletonization and histogram analysis of diffusion tensor magnetic resonance imaging (DT-MRI) data. PSMD was examined in comparison with other brain imaging markers in patients with inherited and sporadic cerebral small vessel disease, and in patients with Alzheimer’s disease and healthy controls (Baykara et al., 2016). PSMD was reported to be associated with processing speed in all samples; the authors reported that it correlated with speed more strongly than other brain imaging markers such as brain volume, volume of white matter hyperintensities, and volume of lacunes.

However, although a special association between PSMD and the cognitive domain of processing speed was emphasised by Baykara et al. (2016), the test that was used to examine processing speed in most samples—the Trail Making Test—is not specific to that domain; Trail Making Test performance is associated with general fluid cognitive ability as well as processing speed (Salthouse, 2011; MacPherson et al., 2017). Moreover, no other cognitive domains were examined by Baykara et al. (2016) for their associations with PSMD. It is therefore not clear whether PSMD correlates with purer processing speed tests, or with processing speed more than it does with other cognitive domains. In addition, there were no formal comparisons undertaken by Baykara et al. (2016) between PSMD’s and other brain imaging biomarkers’ associations with ‘processing speed’ (mostly Trail Making Test performance), nor with other cognitive domains. This means that there was no formal test of whether PSMD correlated better with processing speed—or with other cognitive domains—more strongly than with other structural brain imaging biomarkers. Such a series of tests is necessary before concluding that PSMD is a special neuro-biomarker of the important cognitive domain of processing speed. As Schmidt (2017) explained, all cognitive domains correlate positively together, and, therefore, associations with any one domain might represent an association with general cognitive function rather than a domain-specific relation.

To test whether PSMD has a privileged association with processing speed has the following desiderata, all of which are met in the present study. First, there should be appropriate, relatively-specific tests of processing speed, to test their associations with PSMD. Second, there should be a variety of other cognitive measures, to enable tests of the relation of PSMD to processing speed in comparison with other cognitive domains. Third, PSMD should be compared with other structural brain imaging biomarkers to test whether its association with processing speed and other cognitive domains is especially strong. Here, we run these analyses in a narrow-age sample of mostly healthy, community-dwelling older people: the Lothian Birth Cohort 1936 (LBC1936).

## 2. Materials and Methods

### 2.1 Participants

All participants were members of the LBC1936 (Deary et al., 2007, 2012; Taylor et al., in press). This started (LBC1936, Wave 1) as a sample of 1,091 people who were recruited between 2004 and 2007 at about 70 years of age. All lived independently in the community, were generally healthy, and travelled to a clinical research facility for assessment. Most of them had taken part in the Scottish Mental Survey 1947 at age 11 years. They undertook extensive cognitive, medical, biomarker, psycho-social and other assessments. The assessments were repeated at Wave 2, about three years later at mean age 73 years, with the addition of a detailed structural brain MRI scan (Wardlaw et al., 2011). Of the 886 who returned at Wave 2, over 700 agreed to undertake a brain scan. All variables described below were collected at Wave 2, unless otherwise noted. Ethical approval for the LBC1936 study came from the Multi-Centre Research Ethics Committee for Scotland (MREC/01/0/56; 07/MRE00/58) and the Lothian Research Ethics Committee (LREC/2003/2/29). All participants, who were volunteers and received no financial or other reward, completed a written consent form before any testing took place.

### 2.2 Cognitive tests

The cognitive test data used here are from Wave 2, at mean age 73 years. All participants had already taken the same cognitive test battery at age 70 years, ensuring that they were familiar with the tests.

#### 2.2.1 Processing speed

Because processing speed was the key cognitive variable mentioned in relation with PSMD (Baykara et al., 2016), it is apposite that the LBC1936 has been assessed for processing speed in detail. There were five tests of processing speed: two were paper-and-pencil psychometric tests; two were experimental, reaction time tests; and one was a psychophysical test. The psychometric tests were from the Wechsler Adult Intelligence Test-III^UK^: Digit Symbol and Symbol Search (Wechsler, 1998a). The reaction time tests were simple reaction time (8 practice trials, 20 test trials), and 4-choice reaction time (8 practice trials, 40 test trials), assessed on a stand-alone device that was used in the UK’s Health and Lifestyle Study (Deary et al., 2001). The psychophysical test was inspection time, assessed using a bespoke computer program (Deary et al., 2004). The inspection time task required the participant to indicate which of two briefly-presented parallel vertical lines was longer; no speeded response was required, and only the correctness of the response was recorded. Stimuli were backward-masked. There were 150 trials, with ten trials at each of 15 durations, ranging from 6 ms to 200 ms. Durations were presented at random, using a method of constant stimuli.

#### 2.2.2 Visuospatial ability

This was assessed using Matrix Reasoning and Block Design from the Wechsler Adult Intelligence Test-III^UK^ (Wechsler, 1998a), and Spatial Span (total score of forward and backward) from the Wechsler Memory Scales-III^UK^ (Wechsler, 1998b).

#### 2.2.3 Verbal memory

This was assessed using Verbal Paired Associates (total score) and Logical Memory (immediate and delayed total score) from the Wechsler Memory Scales-III^UK^ (Wechsler, 1998b) and Backward Digit Span from the Wechsler Adult Intelligence Test-III^UK^ (Wechsler, 1998a).

#### 2.2.4 Crystallised ability

This was assessed using the National Adult Reading Test (Nelson and Willison 1991), the Wechsler Test of Adult Reading (Holdnack, 2001), and a phonemic verbal fluency test (using the letters C, F, and L; Deary et al., 2007).

### 2.3 Demographic and health-related variables

The participant’s own occupational social class was based on their most prestigious job prior to retirement, and assessed on a 6-point scale from manual to professional (Office of Population and Census Studies, 1980). Education was assessed as the number of years of full time education. These were assessed at Wave 1. Smoking was assessed at interview and classed as never, ex-, or current. The Mini-Mental State Examination (Folstein et al., 1975) was used as a screen for cognitive pathology, and not used as part of the cognitive test battery. Participants reported if they had a history of hypertension, cardiovascular disease, or diabetes.

### 2.4 Magnetic resonance imaging

All MRI data were acquired using a GE Signa Horizon HDxt 1.5 T clinical scanner (General Electric, Milwaukee, WI, USA) using a self-shielding gradient set with maximum gradient strength of 33 mT/m and an 8-channel phased-array head coil. The full details of the imaging protocol can be found in the LBC1936 imaging protocol paper (Wardlaw et al., 2011). Briefly, the DT-MRI examination consisted of 7 T_2_-weighted (b = 0 s mm^-2^) and sets of diffusion-weighted (b = 1000 s mm^-2^) single-shot spin-echo echo-planar (EP) volumes acquired with diffusion gradients applied in 64 non-collinear directions (Jones et al., 2002). Volumes were acquired in the axial plane, with a field-of-view of 256 × 256 mm, contiguous slice locations, and image matrix and slice thickness designed to give 2 mm isotropic voxels.

The repetition and echo time for each EP volume were 16.5 s and 98 ms respectively. DT-MRI data were converted from DICOM (http://dicom.nema.org) to NIfTI-1 (http://nifti.-nimh.nih.gov/nifti-1) format using the TractoR package (http://www.tractor-mri.org.uk) (Clayden at al., 2011). FSL tools (http://www.fmrib.ox.ac.uk/fsl) (Smith at al., 2004) were then used to extract the brain, remove bulk motion and eddy current induced distortions by registering all subsequent volumes to the first T_2_-weighted EP volume (Jenkinson and Smith, 2001), estimate the water diffusion tensor and calculate parametric maps of mean diffusivity (MD) and fractional anisotropy (FA) from its eigenvalues using DTIFIT (Basser and Pierpaoli, 1996).

#### 2.4.1 Peak width of skeletonized water mean diffusivity

Automatic calculation of PSMD followed the procedure described by Baykara et al. (2016) using the freely-available script they provided (http://www.psmd-marker.com). Briefly, the DT-MRI data were processed using the standard Tract-based Spatial Statistics (TBSS; Smith at al., 2006) pipeline available in FSL with histogram analysis performed on the resulting white matter MD skeletons. First, all participants’ FA volumes were linearly and non-linearly registered to the standard space FMRIB 1 mm FA template. Second, a white matter skeleton was created from the mean of all registered FA volumes. This was achieved by searching for maximum FA values in directions perpendicular to the local tract direction in the mean FA volume. An FA threshold of 0.2 was applied to the mean FA skeleton to exclude predominantly non-white matter voxels. Third, MD volumes were projected onto the mean FA skeleton and further thresholded at an FA value of 0.3 to reduce CSF partial volume contamination. Finally, PSMD was calculated as the difference between the 95^th^ and 5^th^ percentiles of the voxel-based MD values within each subject’s MD skeleton.

#### 2.4.2 Other structural brain imaging variables

The estimation of, respectively, normal-appearing grey and white matter volumes, brain atrophy (intra-cranial volume minus total brain volume), general FA and general MD derived from quantitative tractography, white matter hyperintensity (WMH) volume, and visually-rated perivascular spaces (Potter et al., 2015) from LBC1936 Wave 2 have all been described previously, as, mostly, have their associations with cognitive functions (Royle et al., 2013; Valdes Hernandez at al., 2013; Penke et al., 2012; Booth et al., 2013; Ritchie et al., 2015a,b,c; Aribisala et al., in submission). They are included in the present paper in order to compare their cognitive associations alongside those of PSMD, not as new results in themselves.

### 2.5 Statistical analysis

All analyses were run in the lavaan package for R (Rosseel, 2012). Scores on each multi-test cognitive domain (processing speed, visuospatial ability, verbal memory, and crystallised ability) were derived from separate confirmatory factor analyses assessed using structural equation models based on the analytic sample. Scores on general cognitive ability were derived from a hierarchical confirmatory factor analysis where the score on the general cognitive factor represented the shared variance among the four cognitive domains. Associations between brain imaging parameters and cognitive test scores were conducted using linear regression, with p-values adjusted for the False Discovery Rate (FDR; Benjamini and Hochberg, 1995) as indicated in results tables. To test whether the relation of each cognitive test or domain with PSMD was significantly different from its relation with the other brain-derived variables, we used Williams’s test for differences in dependent correlations (implemented using the psych package in R (Revelle, 2018); treating the standardised regression betas as correlations for this purpose. For each test, we accounted for the dependency (correlation) between the brain variables.

As an additional multivariate analysis, we ran a set of structural equation models in which all eight of the brain measures were entered simultaneously as predictors of the latent cognitive scores. This allowed us to test whether PSMD was incrementally significant beyond the more conventionally-studied micro- and macro-structural brain measures. For the general FA and MD factors, we extracted the factor scores to use as predictors, but still estimated the cognitive factors as latent variables. As with the previous analyses, all variables were corrected for sex and age at the time of testing/scanning. Note that we already reported a similar analysis in this sample (Ritchie et al., 2015b). However, the previous analysis did not include PSMD, and its aim was to discover how much of the variance in cognitive domains could be accounted for with structural brain variables. In the present analysis the aim was to test whether PSMD had an incremental contribution in predicting cognitive variation, in a situation where several structural brain variables were included simultaneously.

## 3. Results

Summary demographic, medical, brain imaging, and cognitive results are shown in Table 1. The total number of subjects who agreed to brain imaging was 731 (388 men), and between 656 and 680 of them provided brain imaging data that were able to be used to compute variables for use in this study. Their mean age was 72.7 years (SD = 0.72). About 47% had a history of hypertension, 27% cardiovascular disease, and 10% diabetes. Their mean Mini-Mental State Examination score was 28.8 (SD = 1.4).

**Table 1.**
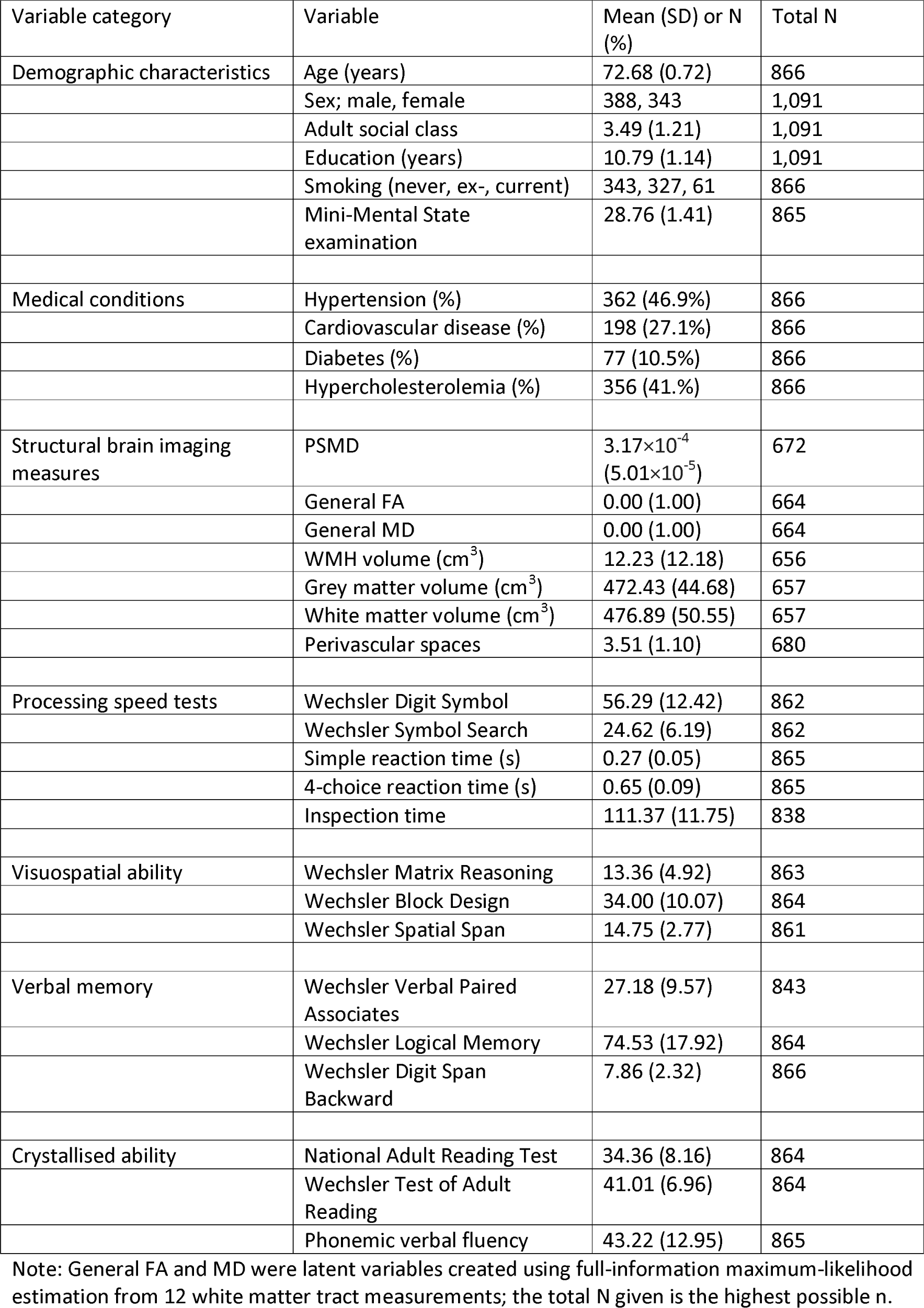
Characteristics of the Lothian Birth Cohort 1936 study sample. Numbers are mean and SD or N and.

| Variable category | Variable | Mean (SD) or N (%) | Total N |
| --- | --- | --- | --- |
| Demographic characteristics | Age (years) | 72.68 (0.72) | 866 |
|  | Sex; male, female | 388, 343 | 1,091 |
|  | Adult social class | 3.49 (1.21) | 1,091 |
|  | Education (years) | 10.79 (1.14) | 1,091 |
|  | Smoking (never, ex-, current) | 343, 327, 61 | 866 |
|  | Mini-Mental State examination | 28.76 (1.41) | 865 |
| Medical conditions | Hypertension (%) | 362 (46.9%) | 866 |
|  | Cardiovascular disease (%) | 198 (27.1%) | 866 |
|  | Diabetes (%) | 77 (10.5%) | 866 |
|  | Hypercholesterolemia (%) | 356 (41.%) | 866 |
| Structural brain imaging measures | PSMD | $3.17 \times 10^{-4}$<br>($5.01 \times 10^{-5}$) | 672 |
|  | General FA | 0.00 (1.00) | 664 |
|  | General MD | 0.00 (1.00) | 664 |
|  | WMH volume (cm<sup>3</sup>) | 12.23 (12.18) | 656 |
|  | Grey matter volume (cm<sup>3</sup>) | 472.43 (44.68) | 657 |
|  | White matter volume (cm<sup>3</sup>) | 476.89 (50.55) | 657 |
|  | Perivascular spaces | 3.51 (1.10) | 680 |
| Processing speed tests | Wechsler Digit Symbol | 56.29 (12.42) | 862 |
|  | Wechsler Symbol Search | 24.62 (6.19) | 862 |
|  | Simple reaction time (s) | 0.27 (0.05) | 865 |
|  | 4-choice reaction time (s) | 0.65 (0.09) | 865 |
|  | Inspection time | 111.37 (11.75) | 838 |
| Visuospatial ability | Wechsler Matrix Reasoning | 13.36 (4.92) | 863 |
|  | Wechsler Block Design | 34.00 (10.07) | 864 |
|  | Wechsler Spatial Span | 14.75 (2.77) | 861 |
| Verbal memory | Wechsler Verbal Paired Associates | 27.18 (9.57) | 843 |
|  | Wechsler Logical Memory | 74.53 (17.92) | 864 |
|  | Wechsler Digit Span Backward | 7.86 (2.32) | 866 |
| Crystallised ability | National Adult Reading Test | 34.36 (8.16) | 864 |
|  | Wechsler Test of Adult Reading | 41.01 (6.96) | 864 |
|  | Phonemic verbal fluency | 43.22 (12.95) | 865 |

Note: General FA and MD were latent variables created using full-information maximum-likelihood estimation from 12 white matter tract measurements; the total N given is the highest possible n.

The age- and sex-adjusted correlations between the brain imaging measures used in this study are shown in Supplementary Table 1. PSMD correlated—in terms of absolute standardized effect sizes—above 0.5 with general FA and MD, and WMH volume, about 0.3 with atrophy and perivascular spaces, and below 0.2 with grey and white matter volumes.

For clarity in understanding the results reported below, all significant associations were in the direction indicating that healthier brains had better performance in processing speed and other cognitive domains. Healthier brains had lower PSMD, higher general FA, lower general MD, lower WMH volume, higher grey and white matter volumes, lower atrophy, and fewer perivascular spaces.

### 3.1 PSMD and other brain imaging variables versus processing speed

We first examined the associations between the five tests of processing speed and PSMD and the other brain imaging measures (Table 2). Higher PSMD (representing less healthy white matter) was significantly associated with worse scores on all five measures of processing speed; the absolute standardised betas were between 0.11 and 0.23 (mean = 0.17). Those with higher PSMD had lower scores on Digit Symbol, Symbol Search, and inspection time, and slower simple and 4-choice reaction times. Therefore, PSMD does correlate significantly, in the expected direction, with these five methodologically-varied tests of processing speed.

**Table 2.** The association between structural brain imaging parameters and individual tests of processing speed.

|  | Wechsler Digit Symbol |  |  | Wechsler Symbol Search |  |  | Simple reaction time |  |  | 4-choice reaction time |  |  | Inspection time |  |  |
| --- | --- | --- | --- | --- | --- | --- | --- | --- | --- | --- | --- | --- | --- | --- | --- |
| | $\beta$ | SE | $p_{adj}$ | $\beta$ | SE | $p_{adj}$ | $\beta$ | SE | $p_{adj}$ | $\beta$ | SE | $p_{adj}$ | $\beta$ | SE | $p_{adj}$ |
| PSMD | -.226 | .037 | $1.01 \times 10^{-08}$ | -.155 | .038 | $5.74 \times 10^{-05}$ | .108 | .037 | .003 | .169 | .037 | $1.04 \times 10^{-05}$ | -.200 | .037 | $3.02 \times 10^{-07}$ |
| General FA | .188 | .041 | $1.70 \times 10^{-05}$ | .119 | .042 | .005 | -.147 | .044 | .001 | -.189 | .042 | $1.70 \times 10^{-05}$ | .172 | .042 | $8.14 \times 10^{-05}$ |
| General MD | -.143* | .041 | .003 | -.092 | .042 | .028 | .121 | .043 | .008 | .113 | .043 | .010 | -.162 | .042 | $5.05 \times 10^{-04}$ |
| WMH volume | -.200 | .037 | $5.55 \times 10^{-07}$ | -.164 | .038 | $3.13 \times 10^{-05}$ | .044 | .037 | .234 | .140 | .037 | $2.12 \times 10^{-04}$ | -.172 | .039 | $2.45 \times 10^{-05}$ |
| Grey matter volume | .259 | .037 | $2.62 \times 10^{-11}$ | .257† | .037 | $3.60 \times 10^{-11}$ | -.047 | .037 | .204 | -.164 | .037 | $1.48 \times 10^{-05}$ | .147 | .039 | $2.05 \times 10^{-04}$ |
| White matter volume | .292 | .036 | $2.55 \times 10^{-14}$ | .257† | .037 | $3.30 \times 10^{-11}$ | -.119 | .037 | .001 | -.215 | .036 | $7.45 \times 10^{-09}$ | .174 | .039 | $1.07 \times 10^{-05}$ |
| Atrophy | .268 | .037 | $4.05 \times 10^{-12}$ | .215 | .038 | $3.16 \times 10^{-08}$ | -.109 | .037 | .003 | -.180 | .037 | $1.42 \times 10^{-06}$ | .220 | .038 | $3.16 \times 10^{-08}$ |
| Perivascular spaces | -.033* | .038 | 0.646 | .005* | .039 | .899 | .059 | .037 | .276 | .005* | .038 | .899 | -.093* | .039 | .085 |

Note. All variables are adjusted for sex and age on the day of testing or MRI scanning. $p_{adj}$ -values are FDR-adjusted within each brain variable (that is, by rows of this table). \* = significantly (absolutely) lower than the $\beta$ for PSMD; † = significantly (absolutely) higher than the $\beta$ for PSMD. Standardised regression coefficients are reported throughout.

Normal-appearing grey and white matter volumes, and brain atrophy had similar associations to PSMD with the five processing speed measures (Table 2); 11 of their 15 betas were larger in effect size (though not significantly larger) than those of PSMD. Associations between WMH volume and the processing speed tests were similar to those of PSMD, though mostly slightly lower. General FA and MD had significant associations with all five of the processing speed measures, with similar betas to those between PSMD and speed measures, though the majority were slightly lower. Perivascular spaces had non-significant associations with the five processing speed measures, and all were notably lower than those with PSMD. We repeated these analyses, omitting participants with a Mini-Mental State Examination score below 24; the results were very similar (Supplementary Table 2).

We tested the effect sizes of the associations between PSMD and individual processing speed tests to find out formally whether they were significantly stronger or weaker than those between other brain imaging variables and the same test. Apart from perivascular spaces, where PSMD always had stronger associations with processing speed measures, PSMD had significantly stronger associations than other brain variables in only one of 30 comparisons (Table 2).

### 3.2 PSMD and other brain imaging variables’ associations with four cognitive domains and general cognitive ability

Next, we examined the possibility that PSMD showed significantly stronger associations with processing speed than with other cognitive domains, and whether they were significantly stronger or weaker associations compared to other brain imaging variables. We ascertained the associations between PSMD and other brain imaging measures and the four cognitive domains (each of which comprised multiple cognitive tests) and general cognitive ability (the variance shared by the four cognitive domains) (Table 3). Higher PSMD was significantly associated, at similar effect sizes, with poorer processing speed (standardised beta = −0.27), visuospatial ability (−0.23), and general cognitive ability (−0.25). The association with verbal memory was significant but lower (−0.17), and the association with crystallised ability was small and non-significant (−0.07).

**Table 3.** The association between structural brain imaging parameters and domains of cognitive ability and general cognitive ability.

|  | Processing speed |  |  | Visuospatial ability |  |  | Verbal memory |  |  | Crystallised ability |  |  | General cognitive ability |  |  |
| --- | --- | --- | --- | --- | --- | --- | --- | --- | --- | --- | --- | --- | --- | --- | --- |
| | $\beta$ | SE | $p_{adj}$ | $\beta$ | SE | $p_{adj}$ | $\beta$ | SE | $p_{adj}$ | $\beta$ | SE | $p_{adj}$ | $\beta$ | SE | $p_{adj}$ |
| PSMD | -.273 | .040 | $4.64 \times 10^{-11}$ | -.235 | .042 | $3.66 \times 10^{-08}$ | -.168 | .045 | $2.37 \times 10^{-04}$ | -.072 | .040 | .070 | -.250 | .041 | $2.57 \times 10^{-09}$ |
| General FA | .238 | .046 | $1.07 \times 10^{-06}$ | .174* | .048 | $4.44 \times 10^{-04}$ | .116 | .050 | .021 | .100 | .043 | .021 | .204 | .047 | $2.89 \times 10^{-05}$ |
| General MD | -.178* | .046 | $5.76 \times 10^{-04}$ | -.163* | .047 | .002 | -.117 | .050 | .025 | .003* | .043 | .951 | -.148* | .047 | .003 |
| WMH volume | -.247 | .042 | $1.39 \times 10^{-08}$ | -.132* | .044 | .004 | -.137 | .046 | .004 | -.048 | .040 | .231 | -.184* | .043 | $4.19 \times 10^{-05}$ |
| Grey matter volume | .315 | .039 | $6.42 \times 10^{-16}$ | .310 | .041 | $4.46 \times 10^{-14}$ | .201 | .044 | $3.97 \times 10^{-06}$ | .253† | .037 | $7.12 \times 10^{-12}$ | .357† | .038 | $3.95 \times 10^{-20}$ |
| White matter volume | .364† | .037 | $3.94 \times 10^{-22}$ | .280 | .042 | $1.64 \times 10^{-11}$ | .175 | .045 | $8.82 \times 10^{-05}$ | .239† | .037 | $1.21 \times 10^{-10}$ | .348 | .038 | $2.45 \times 10^{-19}$ |
| Atrophy | .331 | .039 | $5.89 \times 10^{-22}$ | .277 | .042 | $5.63 \times 10^{-11}$ | .200 | .045 | $1.06 \times 10^{-05}$ | .169† | .038 | $1.33 \times 10^{-05}$ | .320 | .039 | $9.88 \times 10^{-16}$ |
| Perivascular spaces | .038* | .043 | .638 | -.054* | .044 | .638 | -.008* | .045 | .867 | .043 | .040 | .638 | -.020* | .043 | .798 |

Note. All variables are adjusted for sex and age on the day of testing or MRI scanning. $p_{adj}$ -values are FDR-adjusted within each brain variable (that is, by rows of this table). \* = significantly (absolutely) lower than the $\beta$ for PSMD; † = significantly (absolutely) higher than the $\beta$ for PSMD. Standardised regression coefficients are reported throughout.

For all five of the cognitive variables, normal-appearing grey and white matter volumes and brain atrophy had stronger associations than PSMD (Table 3). The associations of WMH volume were somewhat lower than those of PSMD, and all but the association with crystallised ability were significant. General FA and MD had significant associations with all cognitive domains, except between MD and crystallised ability, and the betas were mostly slightly lower than those between PSMD and cognitive domains. Perivascular spaces had no significant associations with any cognitive domain.

We repeated these analyses, omitting participants with a Mini-Mental State Examination score below 24; the results were very similar (Supplementary Table 3). Additional analyses show the associations between the various structural brain imaging measures and the individual tests from the cognitive domains of Visuospatial ability, Memory, and Crystallised ability (Supplementary Tables 4, 5, and 6, respectively).

We tested the effect sizes of the associations between PSMD and each cognitive domain and general cognitive ability to find out formally whether they were significantly stronger or weaker than those between other brain imaging variables and the same cognitive variable. Table 3 shows these results. There were 35 comparisons: in 11 of these, PSMD had stronger associations with cognitive domains and general cognitive ability; and in five of these the PSMD associations were significantly weaker. For processing speed, the cognitive domain of principal interest here, PSMD was a significantly stronger associate than general MD and perivascular spaces, and significantly weaker than white matter volume. Therefore, although PSMD does correlate significantly, in the expected direction, with these cognitive measures, it did not exhibit the largest association among all brain measures in any cognitive domain.

### 3.3 Multivariate models to test PSMD’s incremental contribution to predicting variance in cognitive domains and general cognitive ability

The results of the multivariate models, where all brain variables were entered together to predict variance in cognitive domain scores and general cognitive ability, are shown in Table 4. In these models, PSMD had a statistically significant association with visuospatial ability and with the general cognitive ability factor, but not with processing speed, verbal memory, or crystallized ability. No one brain measure consistently emerged as a statistically significant predictor for each cognitive ability; we note that grey matter volume, white matter volume, and atrophy were all independently significant alongside PSMD in predicting general cognitive ability. The general FA and MD variables were not independently significant in any of the models, indicating that variation in the cognitive abilities was better measured by either the macrostructural indices (except perivascular spaces, which were also not significant in any model), or by PSMD.

**Table 4.** Multivariate models predicting latent cognitive factors from all brain variables, entered simultaneously.

|  | Processing speed |  |  | Visuospatial ability |  |  | Verbal memory |  |  | Crystallised ability |  |  | General cognitive ability |  |  |
| --- | --- | --- | --- | --- | --- | --- | --- | --- | --- | --- | --- | --- | --- | --- | --- |
| | $\beta$ | SE | $p_{adj}$ | $\beta$ | SE | $p_{adj}$ | $\beta$ | SE | $p_{adj}$ | $\beta$ | SE | $p_{adj}$ | $\beta$ | SE | $p_{adj}$ |
| PSMD | 0.093 | 0.060 | 0.240 | **-0.184** | **0.063** | **0.024** | -0.095 | 0.068 | 0.416 | -0.043 | 0.058 | 0.525 | **-0.146** | **0.060** | **0.032** |
| General FA | -0.010 | 0.051 | 0.969 | -0.066 | 0.055 | 0.365 | -0.045 | 0.058 | 0.534 | 0.037 | 0.050 | 0.525 | -0.026 | 0.052 | 0.813 |
| General MD | 0.002 | 0.052 | 0.969 | -0.053 | 0.055 | 0.447 | -0.043 | 0.059 | 0.534 | 0.067 | 0.051 | 0.470 | -0.008 | 0.052 | 0.880 |
| WMH volume | **0.140** | **0.053** | **0.021** | 0.034 | 0.057 | 0.548 | -0.076 | 0.060 | 0.416 | -0.008 | 0.052 | 0.872 | -0.062 | 0.054 | 0.406 |
| Grey matter volume | -0.035 | 0.059 | 0.788 | **0.173** | **0.062** | **0.024** | 0.099 | 0.069 | 0.416 | 0.125 | 0.057 | 0.112 | **0.149** | **0.059** | **0.032** |
| White matter volume | **-0.223** | **0.055** | **$1.83 \times 10^{-04}$** | 0.120 | 0.059 | 0.084 | 0.047 | 0.064 | 0.534 | 0.132 | 0.053 | 0.104 | **0.170** | **0.056** | **0.008** |
| Atrophy | **-0.220** | **0.048** | **$3.69 \times 10^{-05}$** | 0.122 | 0.052 | 0.051 | 0.124 | 0.056 | 0.224 | 0.056 | 0.047 | 0.470 | **0.171** | **0.049** | **0.004** |
| Perivascular spaces | -0.023 | 0.042 | 0.788 | -0.030 | 0.045 | 0.548 | 0.028 | 0.048 | 0.552 | 0.032 | 0.042 | 0.525 | 0.016 | 0.043 | 0.816 |
| $R^2$ | 0.229 | | | 0.159 | | | 0.082 | | | 0.081 | | | 0.214 | | |

Note: Values in bold had significant $p_{adj}$ -values. $p_{adj}$ -values were FDR-adjusted within each multivariate model (i.e. within columns of this table).

## 4. Discussion

In this large sample of community-dwelling older people with a narrow age range around 73 years, we attempted a replication of the previously-reported links between PSMD and processing speed (Baykara et al., 2016). Importantly, we extended the analysis to include necessary comparisons with a variety of more specific processing speed tests, other cognitive domains, and other brain measures. Consistent with the results of the first PSMD-cognitive function report (Baykara et al., 2016), higher PSMD correlated significantly with poorer performance on five methodologically-diverse tests of processing speed: two were psychometric, paper and pencil tests; two were experimental, reaction time tests; and one was a psychophysical test, requiring no fast motor response. Higher PSMD correlated significantly with poorer performance on the latent cognitive domain of processing speed, composed of tests of all three types. However, PSMD was not the exclusive or strongest associate of processing speed: normal-appearing grey and white matter volumes and brain atrophy were often slightly stronger, and WMH volume and tractography-based general FA and MD had slightly lower, but still-significant associations. Moreover, PSMD did not have an exclusive or especially strong association with processing speed compared with other cognitive domains; it correlated at similar levels with visuospatial ability and general cognitive ability, though less strongly with verbal memory and crystallised ability. In multivariate analyses, PSMD contributed predictive power, incremental to that of other structural brain imaging variables, to visuospatial ability and general cognitive ability, but not to processing speed.

If PSMD’s association with processing speed had been higher than those of other structural brain indices, and/or if PSMD has been especially strong in associating with the domain of processing speed by comparison with other cognitive domains, then there would have been strong evidence for arguing that PSMD stands out among brain imaging parameters in being especially associated with this important cognitive domain. Because all cognitive domains are quite strongly correlated, it is important always to test other domains before making a claim that a putative biomarker or any other variable has a special association with one of them; quite often, a variable is, in fact, associated with general cognitive function, which means that it will be associated with most individual cognitive domains. This pitfall (i.e. incorrectly claiming variable X has a particular association with cognitive domain Y, in cases where only domain Y was tested) is common and has been discussed at length (Schmidt, 2017).

Nevertheless, the present study’s results might still be evidence of an important and special association between PSMD and processing speed. Although PSMD did no better than other brain imaging parameters in associating with processing speed, and was also associated with other cognitive domains, it might be argued that, because of its ease of computation and tractability, it is superior to measures that are burdensome to compute and which are more general properties of the brain. However, global measures of GM and WM volume can be obtained from a T1 alone, and their automated measurement can also be undertaken in an automated fashion. That is, both PSMD’s convenience and link to more specific brain biology might be used to argue in its favour; the average magnitude of water molecular diffusion in the centre of common white matter pathways provides information on individual variations in white matter microstructure and the specific methods reduce the likelihood of influences of partial volume effects, rather than simply ‘how much’ of a tissue type an individual possesses. Moreover, the similar association of PSMD with three cognitive domains might not detract from the claim that it is a marker of processing speed. This is because processing speed has long been seen as somewhat special among the cognitive domains. It has been argued that faster processing speed might provide a foundation for better cognitive performance in higher cognitive domains and for less cognitive decline in them (Salthouse, 1996; Verhaeghen, 2014; Ritchie et al., 2014). This is because the tests of processing speed are arguably simpler than those of other cognitive domains, often being referred to as elementary cognitive tests. Thus, if links to underlying biological processes are clearer for PSMD compared with other structural brain markers, and for processing speed compared with other cognitive domains, then it may be argued PSMD is primus inter (apparent) pares for the brain indices examined here. Furthermore, code for performing PSMD is freely available and based on the commonly-used TBSS pipeline making it an accessible biomarker of white matter structure.

The study has the strengths of testing a large and age- and culturally-homogeneous sample, all on the same brain scanner. Processing speed was tested thoroughly, at three levels of description (psychometric, experimental, and psychophysical). The other main cognitive domains that show age-related decline—visuospatial reasoning and memory—and crystallised ability were assessed using multiple, well-validated tests. Also, recognising that all cognitive domains correlate strongly, a general cognitive ability variable, capturing their shared variance, was included. The study has some limitations. The homogeneity of age and cultural background mean that we are not able to generalise beyond those. Neither can we generalise beyond their status as relatively-healthy, community-dwelling individuals; it is possible that samples with a broader range of brain pathology might show stronger brain imaging-cognitive domain associations, and possibly a pattern that brings out more of PSMD’s distinctiveness. The original PSMD-validation study (Baykara et al., 2016) had the advantage of multiple clinical and healthy groups, but its limitations included that most of its samples did not have a relatively-specific test of processing speed and there were insufficient tests of other cognitive domains.

The present study assessed the claim that PSMD is associated with processing speed. We found the predicted association, but it was not exclusive or special with respect to other structural brain indices, or with respect to other cognitive domains. However, PSMD’s specificity, and processing speed’s possible special status among cognitive domains, make the PSMD-processing speed association worth exploring and explaining in a wider range of clinical and non-clinical populations.

## Acknowledgements

The work was undertaken as part of the Cross Council and University of Edinburgh Centre for Cognitive Ageing and Cognitive Epidemiology (CCACE; http://www.ccace.ed.ac.uk). This work was supported by a Research into Ageing programme grant (to I.J.D. and J.M.S.) and the Age UK-funded Disconnected Mind project (http://www.disconnectedmind.ed.ac.uk; to I.J.D., J.M.S. and J.M.W.), with additional funding from the UK Medical Research Council (MRC; to I.J.D., J.M.S., J.M.W., M.E.B., S.J.R. and S.R.C.). J.M.W. is supported by the Scottish Funding Council through the SINAPSE Collaboration (http://www.sinapse.ac.uk). CCACE (MRC MR/K026992/1) is funded by the Biotechnology and Biological Sciences Research Council and MRC. The image acquisition and analysis was performed at the Brain Research Imaging Centre, University of Edinburgh (http://www.bric.ed.ac.uk). The authors thank the research team members who collected, processed, collated and checked cognitive, medical, and brain imaging data. We thank Dr Natalie Royle for her work on some of the non-PSMD brain imaging variables used in the present study.

## Disclosure statement

The authors declare no competing financial interests.

